# Simple and rewireable biomolecular building blocks for DNA machine-learning algorithms

**DOI:** 10.1101/2023.07.20.549967

**Authors:** Ryan C. Lee, Ariel Corsano, Chung Yi Tseng, Leo Y. T. Chou

## Abstract

Deep learning algorithms, such as neural networks, enable the processing of complex datasets with many related variables, and have applications in disease diagnosis, cell profiling, and drug discovery. Beyond its use in electronic computers, neural networks have been implemented using programmable biomolecules such as DNA. This confers unique advantages such as greater portability, ability to operate without electricity, and direct analysis of patterns of biomolecules in solution. Analogous to past bottlenecks in electronic computers, the computing power of DNA-based neural networks is limited by the ability to add more computing units, i.e. neurons. This limitation exists because current architectures require many nucleic acids to model a single neuron. Each addition of a neuron to the network compounds existing problems such as long assembly times, high background signal, and cross-talk between components. Here we test three strategies to solve this limitation and improve the scalability of DNA-based neural networks: (i) enzymatic synthesis to generate high-purity neurons, (ii) spatial patterning of neuron clusters based on their network position, and (iii) encoding neuron connectivity on a universal single-stranded DNA backbone. We show that neurons implemented via these strategies activate quickly, with high signal-to-background ratio, and respond to varying input concentrations and weights. Using this neuron design, we implemented basic neural network motifs such as cascading, fan-in, and fan-out circuits. Since this design is modular, easy to synthesize, and compatible with multiple neural network architectures, we envision it will help scale DNA-based neural networks in a variety of settings. This will enable portable computing power for applications such as portable diagnostics, compact data storage, and autonomous decision making for lab-on-a-chips.

## INTRODUCTION

Biomolecular computers are circuits created from an assortment of biological molecules to mimic electronics. These molecules can range from proteins^1, 2^ to nucleic acids^3–7^, and nanoparticles^8–10^. Compared to the wires and transistors in electronic computers, biomolecules have unique advantages such as their small size, electricity-free operation, and ability to directly analyze other biomolecules present in the solution. These features enable applications such as remote environmental sensing^11^, bio-diagnostics^12, 13^, and data storage^14–16^, without electronics and expensive equipment such as sequencers and microscopes. Biomolecular computers can additionally be integrated into portable devices such as paper^12, 17^ and battery-powered handheld devices^18^. Of the many biomolecule mediums, computers made of nucleic acids such as DNA have achieved the highest complexity, and thus the most power, due to its low cost to prototype, programmability, and ease of design with predictable hybridization kinetics^3, 19, 20^.

Existing DNA-based computers can solve combinatorial^21^, arithmetic^5, 22^, and logical problems^4, 23, 24^. More recently, DNA-based machine learning algorithms, such as support vector machines^25^ and winner-take-all neural networks^13, 25–28^ have been developed to solve more complex pattern recognition and classification problems. Different architectures have been developed to build these DNA computers, including enzyme-free reactions based on toehold-mediated strand-displacement^29^ or exchange^30^, as well as enzymatic reactions such as polymerase-mediated strand displacement synthesis^31, 32^ and the PEN-DNA toolbox^33^. However, despite these advances, DNA computing power still lags behind electronic computers. In general, increasing computing power requires increasing the number of computing units, such as the number of transistors in a processor, neurons in a neural network, or gates in a DNA computer. However, increasing the number of computing units in a DNA-based system also increases spurious signals, or “leak”, due to unintended interactions between supposedly orthogonal DNA sequences in solution^26, 27, 34^. As leak grows with network size, the outcome of the computation becomes noisier and less interpretable. To overcome this bottleneck, we aimed to simplify DNA computing architecture by reducing the number of nucleic acids required to build each component. In doing so, we aim to improve both scalability and usability of DNA computers.

Here we describe a polymerase-actuated DNA computing unit termed “biomolecular neuron”, which serve as rewireable building blocks for neural network algorithms. This scheme combines (i) enzymatic synthesis to encode a greater number of input-output connections on a single DNA strand, (ii) solid-phase immobilization to spatially segregate DNA computing units into network layers, and (iii) universal addressing to enable the assembly of different circuits from the same DNA components. Using this scheme, we demonstrate the ability to build circuits with less nucleic acids. Additionally, we demonstrate the construction of different neural network motifs (e.g. cascading, fan-in, and fan-out circuits) by simply rewiring the connectivity between the same DNA computing units, without any changes to sequence design. Finally, our method generates DNA computing units of longer lengths than is feasible via chemical synthesis, and therefore enables greater input-output connectivity. Altogether, this computing scheme generates computing units from fewer DNA sequences with the capability for more connectivity, and built-in modularity through circuit rewiring. We envision that these advantages will enable scalable DNA computing, such as DNA-based deep learning algorithms, for tackling real-world biological problems.

## RESULTS

### Design and features of biomolecular neurons

In an *in silico* neural network (Figure 1a, left), the simplest computing unit is represented by a circle known as a neuron or a node. Each node receives different pieces of information from upstream nodes (i.e. input), computes this information together using an activation function, and then propagates the result to downstream nodes (i.e. output) (Figure 1b, left). These nodes are arranged into inter-connected layers to form a network mimicking neuronal connections found in the human brain. During the learning (or training) phase, each input (x1, x_2_, x_3_, etc.) is assigned a respective weight (w_1_, w_2_, w_3_, etc.) depending on its contribution to the outcome. Once an optimal weight has been assigned to each connection, the algorithm can be used for its trained purpose.

**Figure 1.**
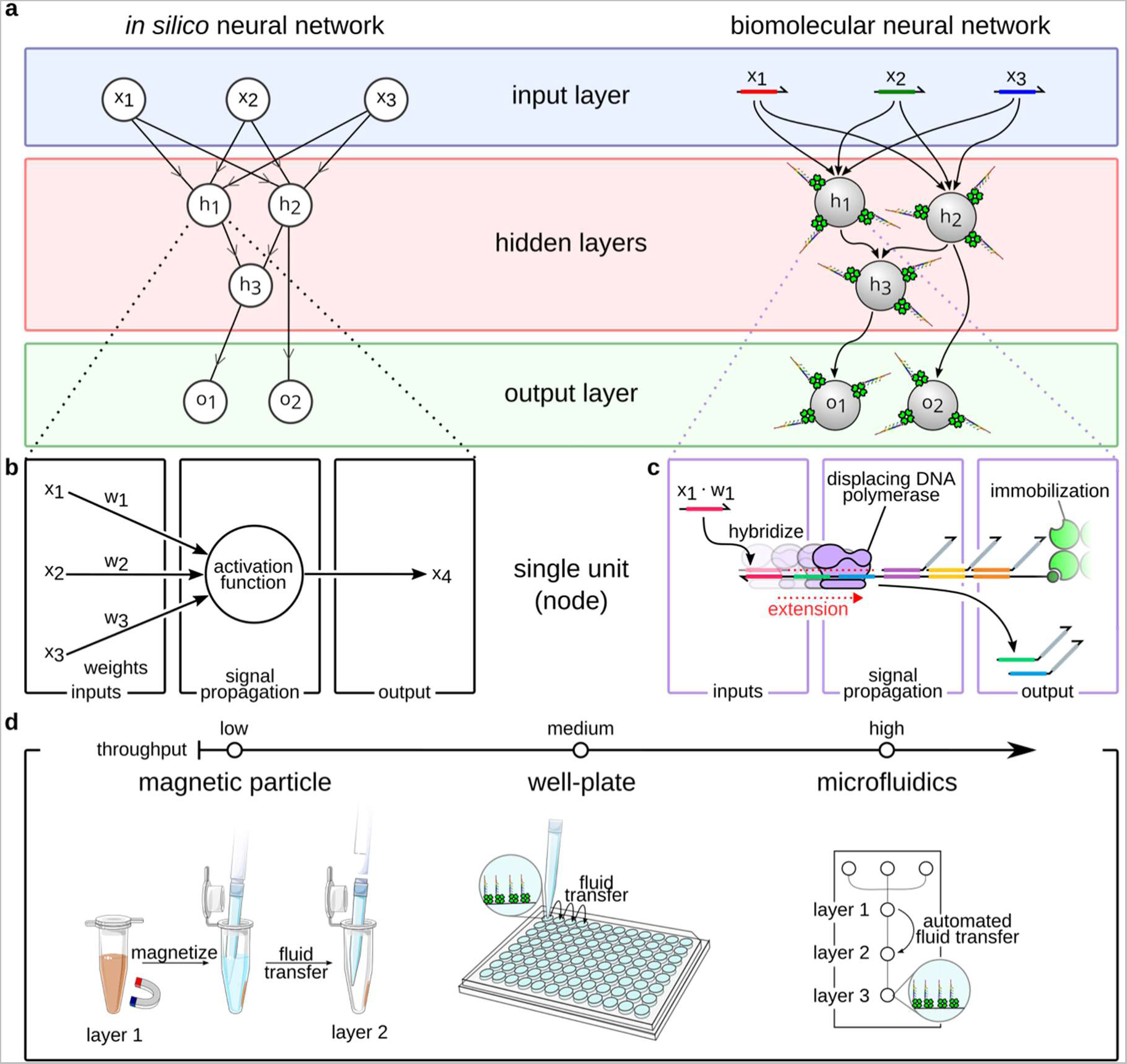
Overview of biomolecular neurons as DNA building blocks for neural networks. **a**, Abstraction of the layers composing an *in silico* neural network (**left**) to a neural network of biomolecular neurons (**right**). Neural networks are composed of an input (x1, x2, and x3) and an output layer (o1, o2), sandwiching two hidden layers (h1 and h2, and h3 respectively). **b,** The structure of a basic computing unit of a neural network, also known as a node. A node receives weighted inputs x1, x2 and x3 with weights w1, w2 and w3 respectively passed through an activation function to produce output x4. **c,** The structure of the proposed biomolecular neuron, separated into its input, signal propagation, and output domains. A ssDNA “backbone” is immobilized onto a solid support, and is equipped with universal addresses allowing for the binding of ssDNA outputs through address-complementary domains. In the presence of a sequence-specific ssDNA input (x1), with binding energy dependent weights (w1), the biomolecular neuron actuates through polymerase-mediated strand displacement synthesis, releasing ssDNA output sequences into the solution. Multiple neurons receive unique input sequences to form a single node of the neural network. **d,** Example workflows for biomolecular neural network implementation from low throughput (left) to high throughput (right). **Left:** Biomolecular neurons are bound to streptavidin coated magnetic particles. Each layer of the neural network is contained in separate tubes, and pulled down with a magnet after reaction before the reaction fluid (containing output ssDNA outputs, enzyme, and other reagents) is transferred to the next layer. This fluid transfer occurs manually with a pipette. **Middle:** Each layer of the biomolecular neural network is contained in a different well of a streptavidin-coated well-plate. Fluid transfer occurs manually with a pipette between layers. **Right:** Neurons of each layer are deposited into separate reaction chambers on a microfluidic device. Fluid transfer is performed by automated fluid pumps. Neural net schematic drawn using publicly available NNsvg tool^46^.

We designed biomolecular neurons to mimic the architecture of *in silico* neurons and form a corresponding biomolecular neural network (Figure 1a, right). Each biomolecular neuron is represented by a long single-stranded DNA (ssDNA) backbone, while the inputs and outputs are represented by shorter ssDNA oligonucleotides. The ssDNA backbone consists of multiple 20-nucleotide (nt) domains that serve as universal addresses for input and output binding. Each output sequence has two domains, the first binds to an address on the backbone through complementary base-pairing, and the second is a 3’ single-stranded overhang which is complementary to the input domain of another neuron. Thus, the same neuron can be swapped with different output sequences to redefine its connections between other neurons, without modifications to the backbone (i.e. rewiring). When not in use, addresses can be filled using output sequences with non-interactive 3’ terminus instead. Using this modular swapping, we can rewire predesigned neuron backbones into a variety of circuit architectures. Furthermore, this design allows better circuit scalability, since each additional neuron requires the design of only one new orthogonal domain (i.e. the input domain).

The biomolecular neuron actuates through polymerase-mediated strand displacement synthesis^22, 31, 32^ (Figure 1b, right). When a free-floating ssDNA input sequence from solution binds to its complementary domain on a biomolecular neuron, it serves as a primer for extension by a strand-displacing polymerase. Primer extension along the backbone in turn displaces the bound output strands into solution. To separate these output strands from the dsDNA extension by-product, we anchored the neurons onto a solid surface. After a short reaction period, the solution containing the polymerases and accumulated outputs is withdrawn and transferred into a new container with the next layer of unreacted neurons, while the by-products remain in the old container. This fluid transfer step can be accomplished using different container types with varying accessibility, setup time, and throughput (Figure 1d). For example, a manual reaction uses biotinylated neurons anchored onto streptavidin-coated magnetic particles, which can be pulled down while the fluids are transferred via pipetting. While this approach has low throughput, it has the advantage of being cheap and easily accessible. Alternatively, the use of streptavidin-coated well-plates remove the need to magnetize samples between reactions and enable serial reading of samples for higher throughput. Finally, integration with microfluidics would enable fully automated fluid handling for high throughput and allows multiple parallel reactions to occur simultaneously. Together, our proposed biomolecular neuron provides a system for biomolecular computing which can be used with different levels of lab infrastructure.

### Biomolecular neuron synthesis and validation

To synthesize the long ssDNA that forms the backbone of our biomolecular neuron, we use polymerase chain reaction (PCR) rather than chemical synthesis (Figure 2a). This allows us to generate longer backbones for encoding a greater number of input-output connections on a single neuron. It also ensures full-length sequences to circumvent leak induced by chemical synthesis defects, and removes the need for laborious purification steps such as polyacrylamide gel electrophoresis (PAGE) or high-performance liquid chromatography (HPLC). The PCR reaction uses a pair of chemically modified primers. The forward primer contains a 5’ phosphate that enables downstream digestion of the dsDNA amplicon into ssDNA, and a 20-nt 5’ overhang encoding the variable input-complementary domain (x_n_) that is appended onto the neuron during PCR. The reverse primer contains a 5’ biotin modification for solid-phase immobilization. Following PCR, dsDNA amplicons are converted into ssDNA via lambda exonuclease digestion (Supplementary Figure 1). Here, we note that we tested other ssDNA isolation strategies before settling on enzymatic digestion. For example, we tested a NaOH denaturation step post-immobilization seen in published SELEX protocols^35^. However, similar to other groups, we found that this approach damaged solid supports and resulted in large batch-to-batch variability of total ssDNA (Supplementary Figure 2)^35–37^. Following ssDNA synthesis, the ssDNA backbone is hybridized with outputs containing the address-complementary domains to form neurons. Finally, the neurons are immobilized to the solid support and excess ssDNA is washed away (Figure 2a). Together, this digestion and immobilization protocol allows fast and cheap neuron synthesis, bypassing laborious native PAGE purification steps traditionally used to remove stoichiometry errors^35, 36^.

**Figure 2.**
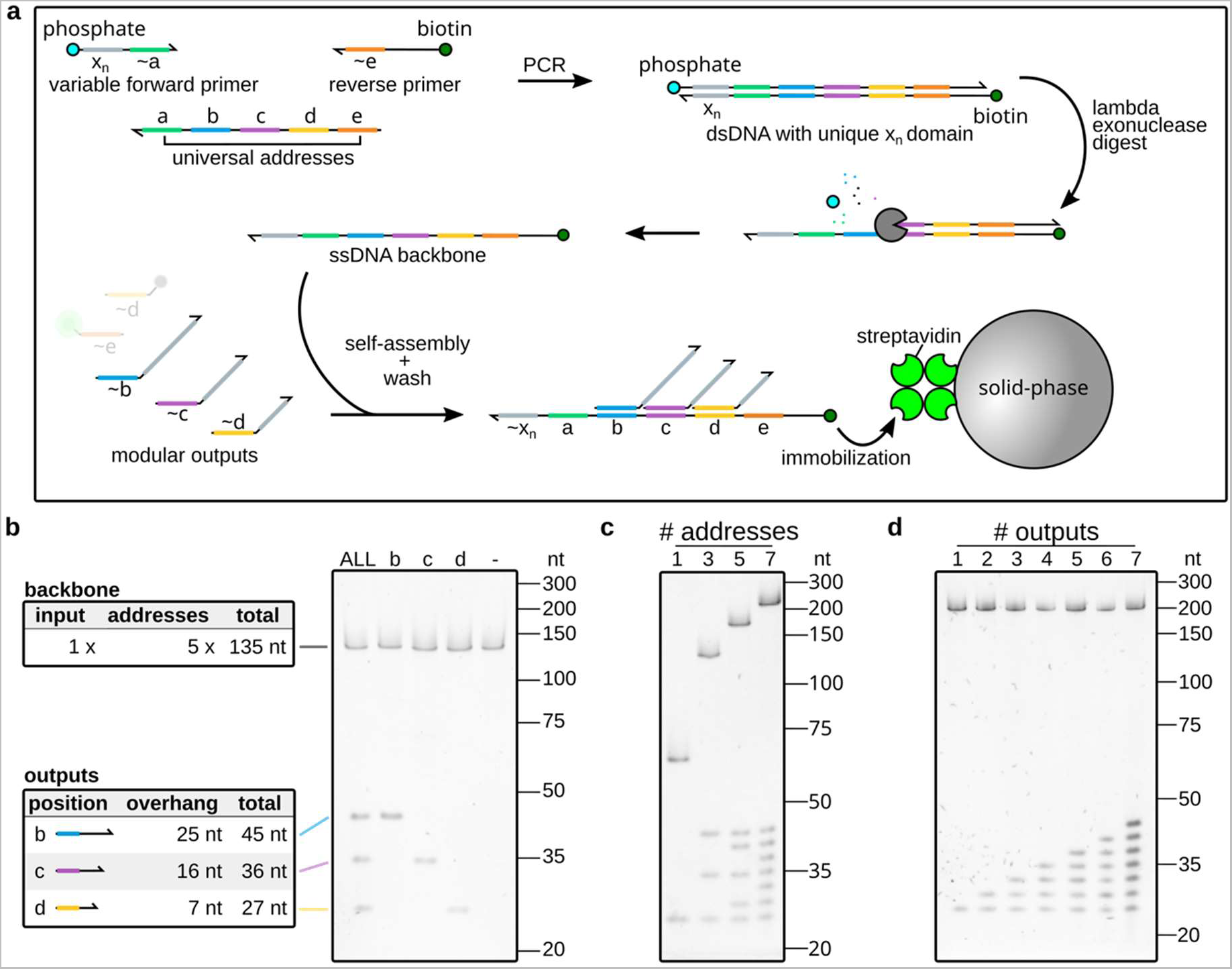
Workflow for modular synthesis of biomolecular neurons. **a**, Workflow for modular neuron assembly. A 5’ phosphate primer with input (X_n_) overhang and a 5’ biotin primer are used to amplify a template with universal addresses (a, b, c, d, e) through PCR. Lambda exonuclease digests the 5’ phosphate tagged strand, leaving a ssDNA backbone. Outputs with different functional overhangs are then modularly annealed (through address hybridization) before immobilization to a solid support. Liquid phase washes remove any unhybridized nucleic acids. **b**, Addresses b, c, and d were labelled separately, and together on a 135nt 5-address backbone. Samples were loaded and stained on denaturing polyacrylamide gel (dPAGE) to show labelling of addresses with different “output” oligonucleotides with varying overhang lengths serving as barcodes for strand identity. Addresses a and e left empty for other potential labelling strategies. **c,** dPAGE shows increase in neuron connectivity by increasing number of addresses on backbone. 1, 3, 5, and 7 addresses were labelled on backbones of different lengths. **d,** dPAGE shows modular and programmed labelling of addresses. 7 addresses were labelled in increasing numbers to demonstrate the ability to modularly swap output oligonucleotides. All sequences used in this figure are available in Supplementary Table 1.

To validate our synthesis protocol, we aimed to verify the ability to (i) produce different neuron backbones from a single dsDNA template using PCR, and (ii) subsequent output hybridization on the backbone using address-complementarity. To do this, we designed three forward primers each containing a unique 20-nt input domain, and used PCR to append them to a 112-nt DNA template containing 5 universal addresses. We characterized the backbones using denaturing PAGE to verify that in all cases, amplicon and ssDNA of the expected size was generated (Supplementary Figure 1). To demonstrate output functionalization, we labeled one of the ssDNA backbones on the second, third, and fourth addresses (i.e. domains b, c, and d in Figure 2b) using output oligonucleotides containing different overhang lengths as barcodes for their identities (Figure 2b). This assay shows that each address could bind its respective output oligonucleotide without any positional dependencies. Addresses a and e were initially left unhybridized for other potential functionalization processes, but also exhibited similar binding capabilities (Supplementary Figure 3). We further quantified the variability of labelling using fluorescently-tagged outputs to verify that biomolecular neurons assemble properly without sequence-dependent effects (Supplementary Figure 4). We used this protocol to create the three different 5-address backbones used in the remainder of this work.

**Figure 3.**
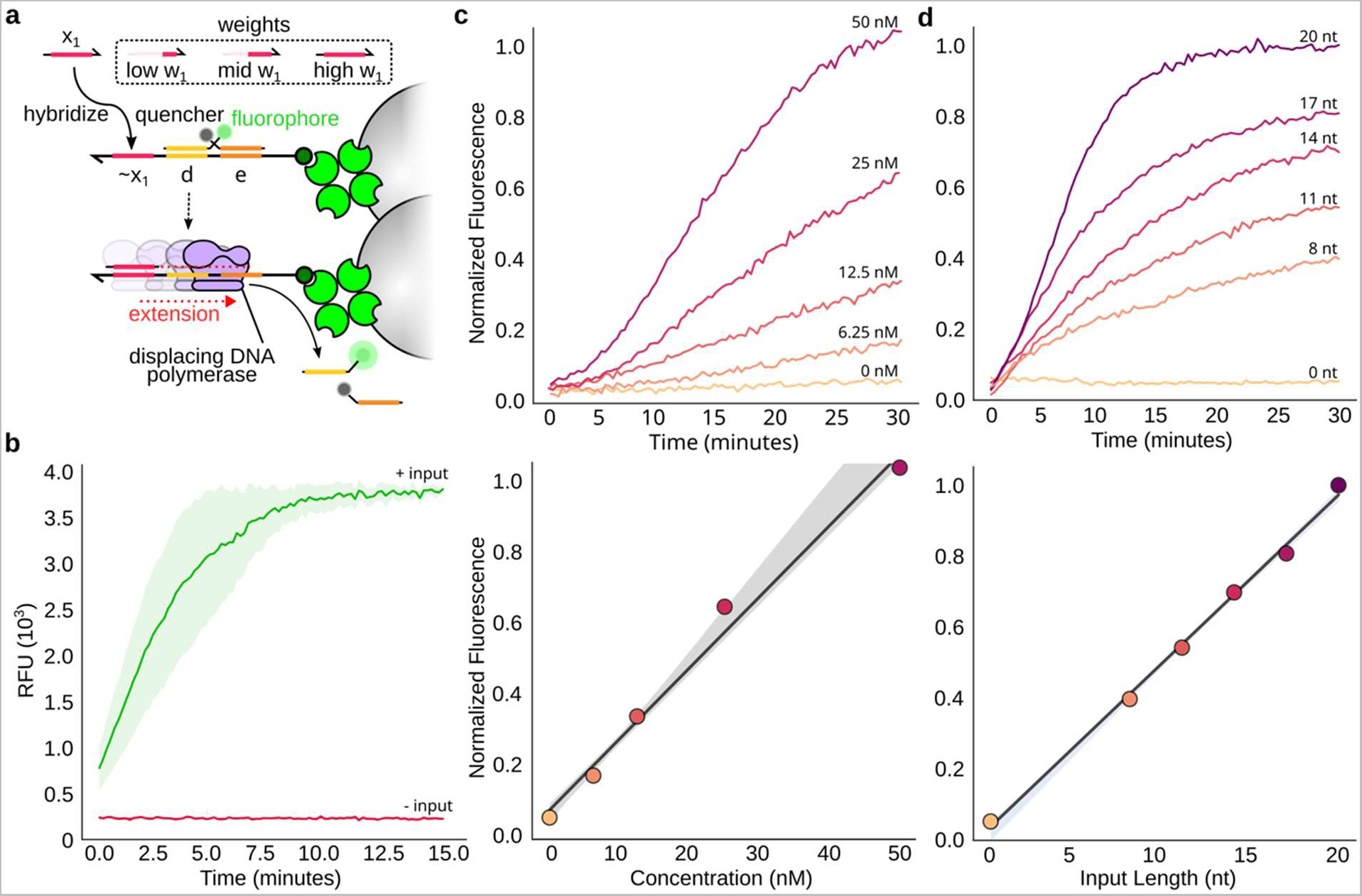
Biomolecular neurons enable modelling of values and weights. **a**, Schematic of a neuron with a fluorophore/quencher pair for kinetics measurements. A displacing DNA polymerase extends a bound input primer to displace the fluorophore/quencher pair, leading to fluorescence recovery. Input DNA can be varied in length to determine the weight (w1, w2, w3, etc.) of the connection. **b**, Kinetic curve of neuron activation using fluorophore/quencher pair, with and without input DNA at 100 nM. Results show signal-to-background ratio of 10.33 ± 0.7 (SEM), n=3. Shaded area indicates SEM. **c, Top:** Kinetic curve of neuron response to different concentrations of input in solution ranging from 0 to 50 nM. **Bottom:** scatter plot of neuron response endpoints to concentration. Shaded area indicates 68% confidence interval. **d, Top:** Kinetic curve of neuron response to different weighted inputs by varying hybridization length from 20 nt to 8 nt. **Bottom:** Scatter plot of neuron response endpoints to input length. Shaded area indicates 68% confidence interval. Fluorescence data in this figure was collected with streptavidin-coated magnetic beads, and data from panels c & d was normalized to a positive control with a large excess of input (100 nM), n=3.

**Figure 4.**
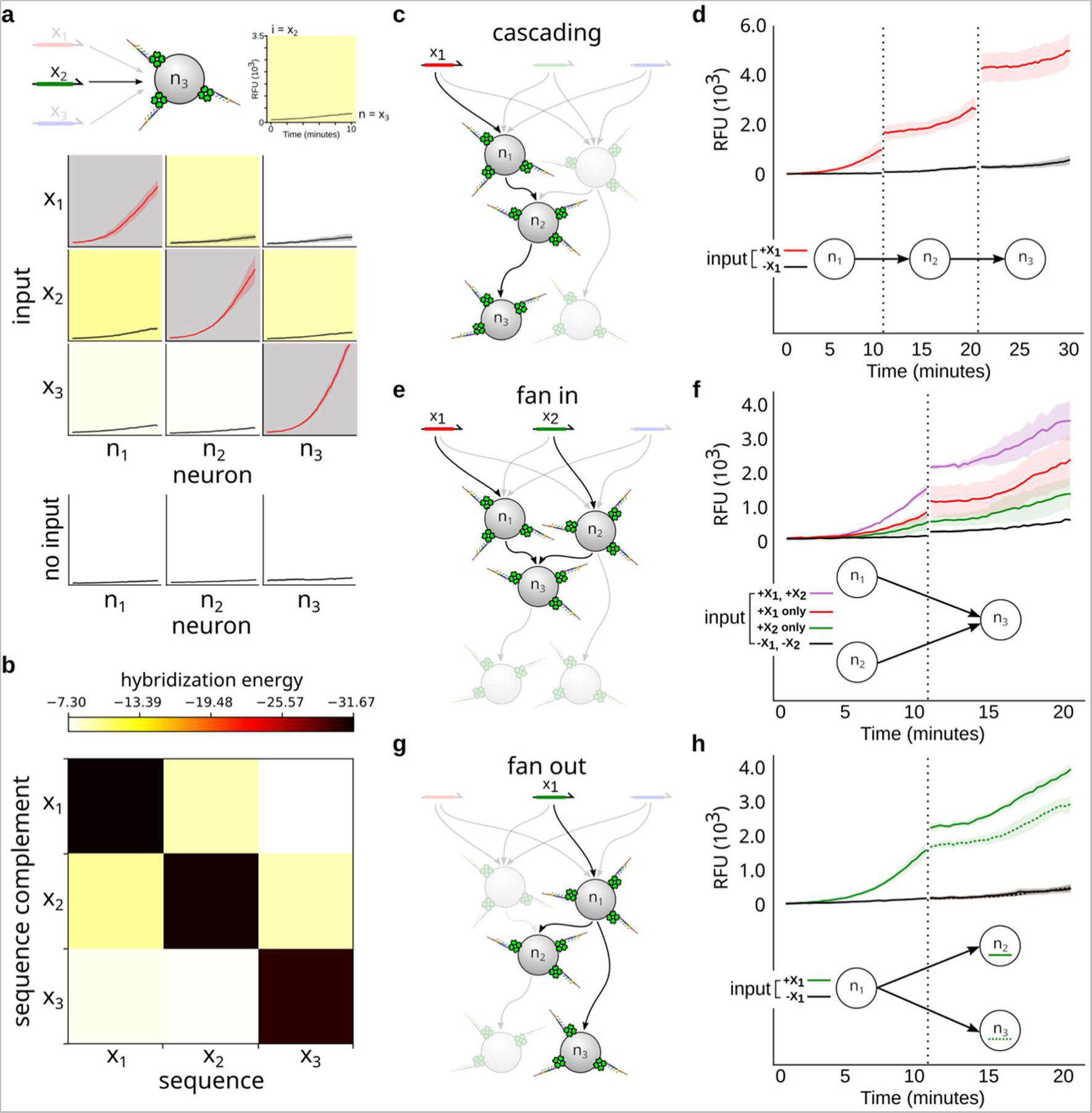
Biomolecular neurons can be rewired into various neural network motifs. **a**, A heatmap of the activation of three neurons against their input sequences. Activation occurs only in the presence of the neuron-specific input, highlighted by the grey background. Shaded areas indicate SEM, n=3. **b,** NuPACK analysis of hybridizing energies (kCal/mol) of input and input-complement domains, used in a. **c,** Abstraction of a 3-neuron cascade as a subset of a biomolecular neural network, where n1 activates n2, which in turn activates n3. **d,** Kinetic trace of implementation of a 3-neuron cascade in biomolecules with manual fluid transfer indicated by dotted lines after 10 minutes of reaction. +x1 condition (+ input) indicated in red, -x1 condition (-input) indicated in black. **e,** Abstraction of a fan-in circuit as a subset of a biomolecular neural network, where n1 and n2 converge to both activate n3. **f,** kinetic trace of implementation of a fan-in circuit using biomolecular neurons in the presence of different combinations of two inputs (+x1/+x2, +x1/-x2, -x1/+x2, -x1/-x2). Inputs x1 and x2 were added at 30 nM and 20 nM respectively. Results show activation of n3 by either n1 or n2, and summation when both inputs are present. **g,** Abstraction of a fan-out circuit as a subset of a biomolecular neural network, where n1 activates both n2 and n3. **h,** Implementation of a fan-out circuit using biomolecular neurons with and without the input (x1). Activation of the two downstream neurons, n2 and n3 are denoted by the solid and dotted lines respectively. Fluorescence data in this figure was collected with streptavidin-coated well-plates (n=3) and not normalized. Shaded areas of the curves indicate SEM. All sequences used in this figure are available in Supplementary Table 2.

We envisioned that different applications of biomolecular neural networks would require different network sizes and computing power. Therefore, we synthesized 3 other backbones with differing numbers of address domains to fulfill this need for more inter-neuronal connectivity. In addition to the backbone with 5 addresses (a, b, c, d, e) (Figure 2b), we synthesized shorter (3– address) and longer (7– and 9–address) backbones to allow for differing connectivity. We again verified the labelling of these new backbones using denaturing PAGE (Figure 2c). Finally, we show that the 9-address backbone can be labelled at any one of its addresses in user-defined combinations (Figure 2d). Together, these results demonstrate flexibility in neuron length and labelling for neural networks of different sizes.

### Neuron actuation is fast and responds to varying input concentrations and weights

Following neuron synthesis, we next investigated its polymerase-mediated actuation. Our reaction used Bst 3.0 polymerase and included a single-stranded binding protein (SSB) from *E. coli*, which we found to reduce polymerase mis-priming (Supplementary Figure 5), similar to prior studies^38^. For initial testing of neuron activation, we used a neuron containing one input-complementary domain and two address domains (d and e) (Figure 3a). We functionalized the neuron with output oligonucleotides labeled with a 3’ BHQ2 on address d and a 5’ FAM on address e. In this way, primer extension by the polymerase would displace the fluorophore- and quencher-labeled oligonucleotides, resulting in an increase in fluorescence signal. Using this fluorophore/quencher scheme, we found that our reaction environment activates neurons maximally within 15 minutes (Figure 3b), and reaches an average 10-fold (10.33 ± 0.7, mean ± SEM, n=3) signal-to-background ratio, similar to other blunt-end fluorophore/quencher designs^22^. Thus, we used this reporting scheme in the remainder of this work to characterize neuron activation kinetics. We note that our reactions were performed at 40℃, lower than the 50-72℃ suggested for optimal Bst 3.0 polymerase activity^32^. This was to reduce the length of the DNA domains needed for circuit building. Nevertheless, we anticipate that faster kinetics could be achieved by operating at the optimal temperature of the polymerase.

**Figure 5.**
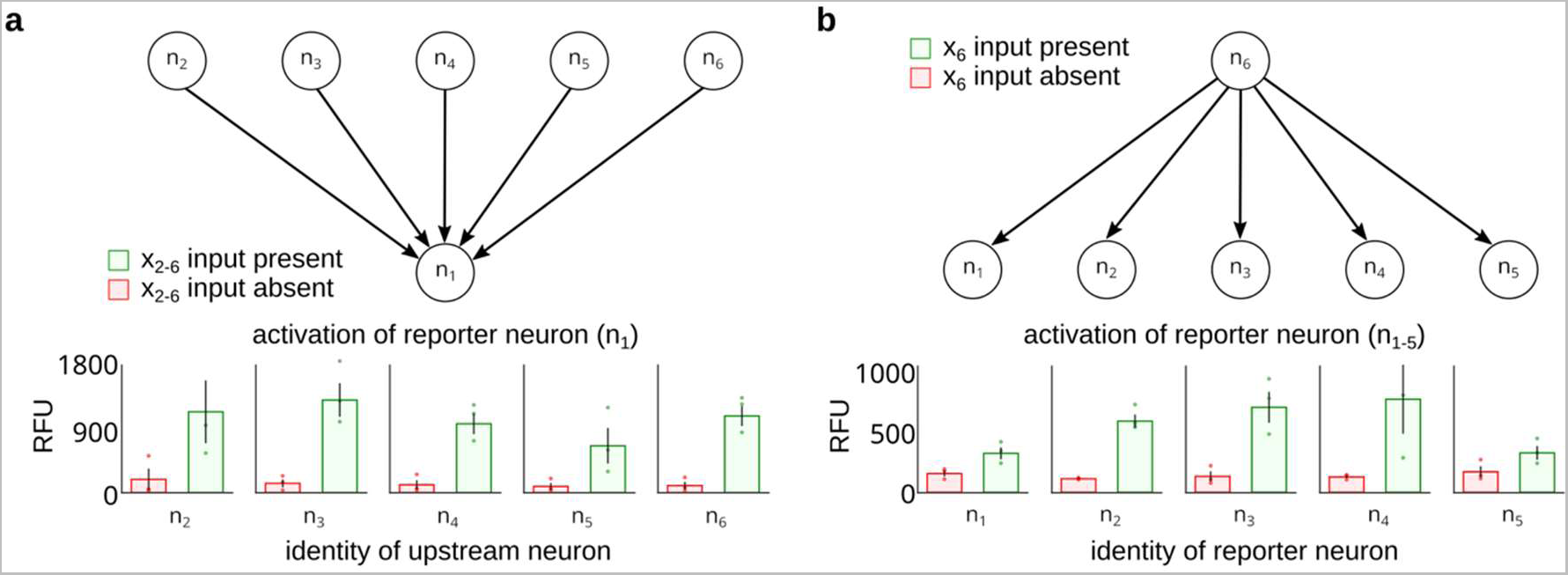
Biomolecular neurons enable increased connectivity of fan in and fan out circuits. **a**, 5-input fan-in circuit created from rewiring of six (6) biomolecular neurons. Neurons n2 to n6 are wired via universal backbone with x1 outputs which activate n1, and n1 is wired with the fluorophore/quencher scheme. Fluorescence of the downstream reporting neuron, n1, is recorded when each upstream neuron is activated by their respective input (ie. n2 is in the presence of x2), and their fluorescence intensities at 10 minutes are plotted in green. Similarly, fluorescent signals of input-absent conditions are plotted in red. Results show activation of n1 only when the input of the upstream neuron is present. **b,** 5-output fan-out circuit created from rewiring of six (6) biomolecular neurons. Each of the five addresses of neuron n6 is assigned a different output strand, activating neurons n1 through n5, and neurons n1 through n5 are wired with the fluorophore/quencher scheme. Fluorescence of the downstream reporter neurons (n1 through n5) are measured in either the presence or absence of the input x6. Fluorescence intensities are recorded at 10 minutes and input present conditions are plotted in green, while input absence conditions plotted in red. Fluorescence data in this figure was collected using streptavidin-coated well-plates (n=3) and not normalized. Error bars indicate SEM. All additional sequences used in this figure are available in Supplementary Table 3.

In artificial neural networks, inputs can take on a continuous set of values. Similarly, we aimed to demonstrate that biomolecular neurons can respond to varying input values represented as the concentration of the input oligonucleotide. To do this, we reacted the fluorophore/quencher labeled neuron with input concentrations starting at 6.25 nM, and was found to activate maximally at 50 nM (Figure 3c, top). We found that increasing concentrations resulted in higher endpoints and faster kinetics, and found a linear relationship between neuron response and concentration in this range (Figure 3c, bottom). In addition, we found that concentrations above this range (>100 nM) resulted in no signal increases above the 50 nM condition, but increased kinetics instead (Supplementary figure 6). From these results, we concluded that numerical values could successfully be encoded by the concentration of nucleic acids, under 50 nM.

We next aimed to demonstrate weighted connections between neurons by manipulating the length of the hybridizing input domain, analogous to adjusting toehold displacement kinetics by manipulating toehold length^34^. To do this, we tested oligonucleotide strands with varying lengths as inputs, from 7 nt to 20 nt, and plotted their kinetic traces (Figure 3d, bottom). As expected, strands with longer inputs domains, and thus stronger hybridization energies, reached higher fluorescent signal than those with shorter domains. We found a linear relationship between input length and signal amplitude This relationship remained true when plotted as a function of hybridization energy instead of domain length (Supplementary Figure 7), suggesting that one can implement weighted connections via DNA hybridization simulations such as NuPACK^20^. Altogether, these results indicate that biomolecular neurons can activate quickly in minutes, and respond to inputs of varying concentrations representing signal value, as well as varying lengths representing signal weight.

### Neurons can operate orthogonally and assemble into networks

To demonstrate that neurons are activated specifically by their cognate inputs, we designed three neurons each activated by a different input sequence. We then introduced a different input to each neuron separately, and compared their specific and non-specific activation levels. We found that neurons activated in response to their specific input with an average of 13.28-fold signal-to-background (± 2.57, SEM), compared to 1.75-fold (± 0.29, SEM) for non-specific inputs (Figure 4a). Furthermore, we found that the resulting neuron activation could be predicted by the hybridization energy of the input and the input-complement domain in NuPACK (Figure 4b). These results demonstrate the ability of our design process to produce highly orthogonal sequences with predictable behaviours for scaling up computing power.

Another feature for scalability in neural network architecture is the ability to form signal cascades. To test the ability of our neurons to form a 3-layer cascade (Figure 4c), we used PCR to synthesize three neurons, each with one unique input-complementary domain and five universal addresses. We connected these neurons serially, i.e. neuron 1 (n_1_) activates neuron 2 (n_2_), which then activates neuron 3 (n_3_), by functionalizing three of the addresses with the corresponding output oligonucleotides. The remaining two adjacent domains were labeled with BHQ2/FAM for fluorescence reporting. We anchored these neurons onto streptavidin-coated well-plates. We reacted the well containing n_1_ with enzymes and reagents for 10 minutes, and transferred the resulting supernatant to the next layer (n_2_). We repeated this cascade through three neurons. Figure 4d shows the raw kinetic traces of each neuron, both when the input is present (+x_1_ condition) and absent (-x_1_ condition). Here, we observed stepwise accumulation of fluorescence signals at each layer of the cascade, denoted by the dotted lines. Furthermore, signal accumulation was specific to the +x_1_ condition and was otherwise low in the -x_1_ condition. Interestingly, we observed unexplained jumps in fluorescence after fluid transfer, seen by the gaps in the red curve. We hypothesize that many factors may contribute to these jumps, such as slow manual fluid transfers, gradual adsorption of proteins to well-plate material, and evaporation of the reaction fluid. Despite the unexpected gaps in data, the 10-fold signal-to-background of single-neuron reactions was retained over three cascades without normalization. With these results, we demonstrate the ability to propagate signals through at least three layers, the minimal depth required to build a deep learning neural network^39, 40^.

To demonstrate the ability for neurons to receive multiple different input signals, we designed a fan-in circuit consisting of two neurons in the first layer converging into one neuron in the second layer (Figure 4e). Using the same neuron backbones as the ones used to build the three-layer cascade, we rewired n_1_ and n_2_ to both cascade into n_3_ instead. Neurons n_1_ and n_2_ were both anchored in the first well, while n_3_ was anchored in the second well. Here we add combinations of the two inputs (+x_1_/+x_2_, +x_1_/-x_2_, -x_1_/+x_2_, -x_1_/-x_2_), each denoted by a different coloured trace in the plot, to demonstrate that activation of either upstream neuron (n_1_, n_2_) results in downstream activation of n_3_ (Figure 4f). Furthermore, activation of both first-layer neurons results in summation of the signal, while no activation of the first layer retains a low background signal. With this result, we demonstrate orthogonal pooling of neurons into the same container to form a layer, as well as fan-in connectivity between neurons of different layers.

Next, we aimed to demonstrate the ability of a single neuron to activate multiple neurons. To do this, we rewired the neurons into a fan-out circuit, where a single first-layer neuron (n_1_), activates two neurons in a downstream layer (n_2_, n_3_). Neurons in the second layer were split into different wells to differentiate their activation using only one fluorescent colour. Here we show that a connection of the activated neuron (n_1_) to either downstream neuron (n_2_, n_3_) results in propagation of signal (Figure 4e) denoted by the dotted or solid line. Altogether, we demonstrate the design of highly orthogonal neurons with room for future scalability, and modular rewiring of these neurons into different neural network motifs without changes in sequence design.

### Expanding the connectivity of biomolecular neurons

Finally, we aimed to demonstrate that neuron connectivity was not limited to only two simultaneous connections. To do this, we designed and synthesized three additional neuron backbones, n_4_ through n_6_, using the same five-address backbone. We wired the new total of six neurons into a 5-input fan-in circuit, where neurons n_2_ through n_6_ all converge to activate a reporter neuron n_1_ (Figure 5a). In this design, neurons of the first layer, n_2_ through n_6_, are outfitted with n_1_-specific output sequences on addresses a, b, and c, with the remaining two addresses d and e occupied by non-interactive outputs. Meanwhile, the reporting neuron n_1_ contains the fluorophore/quencher pair on addresses d and e, and non-specific outputs filling addresses a, b and c. In this way, activation of any first-layer neuron would cause sequential activation of the reporter neuron and result in a measurable fluorescent signal. To setup the circuit, we anchored the 5 neurons of the first layer into 5 separate wells of a streptavidin-coated 384-well plate, and then anchored 5 copies of the reporter neuron into 5 other wells. We allowed the first layer to react with their respective inputs (x_2_ - x_6_) for 10 minutes, before transferring the reaction fluid to the reporter neurons. We plotted the resulting fluorescent signals from the reporter neurons after 10 minutes, with input-present conditions in green, and input-absent conditions in red. From these results, we demonstrate that five neurons can be successfully wired to interact with a single neuron, shown by the high fluorescent signal in input-present conditions, while maintaining a low background signal in input-absent conditions.

To demonstrate that all five addresses of the existing neurons could make outbound connections, we re-wired neurons n_1_ through n_6_ into a fan-out configuration, where n_6_ is used to activate neurons n_1_ through n_5_ (Figure 5b). In this design, each address of n_6_ is occupied by an output strand corresponding to a different neuron, while the remaining neurons are outfitted with non-specific outputs and a fluorophore/quencher pair. Using the same streptavidin well-plate experimental setup, we recorded the 10-minute fluorescence signals of the reporter neurons with and without the n_6_-specific input, x_6_, plotted in green and red respectively. These results show that an initial activation of the first-layer neuron n_6_ results in subsequent activation of any of the five downstream neurons, and lack of activation retains the low background signal we expected. We note that compared to the fan-in circuit, the fan-out circuit results in lower signal-to-background ratios. We attributed this to the lack of signal amplification in the latter circuit as all addresses were used to make outbound connections to a different neuron. From these results, we envision that higher degrees of neuron interconnectivity may be possible by increasing the number of addresses on the neuron backbone, allowing for neural networks with greater layer width without exponential increases in number of unique nucleic acid sequences.

## CONCLUSION

In this work, we present biomolecular neurons as a modular building block for nucleic-acid neural networks. By combining enzymatic synthesis, solid-phase immobilization, and universal addresses, we enable accessible, low-cost, and low-labour DNA neural network circuits with minimal DNA strands. Compared to other computing architectures such as toehold-mediated displacement, biomolecular neurons are simpler to implement because of shorter assembly times and fewer unique components. For example, a fan-in circuit with weighted inputs implemented with toehold mediated strand displacement requires 22 DNA sequences and 14 unique components^6^. In comparison, a similar implementation in biomolecular neurons would require a bare minimum of 6 DNA sequences and 3 unique components. Moreover, digital toehold mediated strand displacement circuits require additional gates to encode values (such as in binary), while in an analog implementation of polymerase-mediated strand displacement, these features are naturally built into tunable properties of nucleic acids such as concentration and binding energy of the input sequence. Our system expands on existing polymerase-mediated strand displacement systems by introducing addressed connections; this allows for building different circuits without re-design and re-synthesis of sequences. Finally, a surface-based DNA computing approach has a unique feature; computation at each layer is synchronized to the timing of fluid transfer. We argue that the simpler assembly process, rewiring capability, and synchronized computation of biomolecular neurons will allow for better scalability when developing neural network models with nucleic acids.

Despite the many advantages, we have not yet completely realized our goal of maximizing nucleic acid circuit scalability. We found that cascades through multiple layers resulted in a small, but noticeable level of leak, which may limit the number of sequential layers in a neural network, known as depth. We hypothesize that this leak may be due to a few possible factors, such as biotin-streptavidin off-rate, change in reaction environment due to protein adsorption or fluid evaporation, or undesired exonuclease activity from enzymatic synthesis. We aim to investigate these factors in future work to fully enable networks with deep connections. Additionally, while we demonstrate low- and medium-throughput systems through manual pipetting, we aim to automate this using microfluidics^41–43^ in future works, including designs which do not require peripherals^44^. Finally, techniques such as microarray printing and lithographic patterning can be used to minimize the footprint of the biomolecular neural network. We envision that future work on surface-based DNA computing with our biomolecular neurons can be sufficiently scaled to implement large deep learning networks.

## METHODS

### Sequence design

Sequences in this work were designed using the Python implementation of NuPACK 4.0^14^. To generate these sequences, we designed a small 40-member library of orthogonal addresses, each 20 nucleotides long. Constraints were placed on these addresses such that they had a GC content of 40-60%, and did not contain repeats of 3 or more nucleotides, nor did they contain significant secondary structure (average ΔG = -7.69) at 40℃. Furthermore, we asserted that these addresses do not form any hairpin structures, regardless of binding strength, to guarantee that no polymerase-mediated self-priming could occur. To create the backbone sequences, we appended these (1 input domain + 3, 5, 7, and 9) addresses with 3T spacers in an order resulting in the lowest secondary structure free energy. ssDNA outputs were designed by appending an output sequence of interest (a complement to an input domain) to the complement of an address domain with a 3T spacer. All sequences used in this work are available in the supplementary material (Table S1).

### PCR amplification of biomolecular neuron amplicons

All oligonucleotide sequences were purchased from IDT at various scales (25 nmol to 100 nmol depending on modifications) and purified with standard desalting. 112-nt template was ordered as a ultramer at 4 nmol scale. No further PAGE purification was performed. All sequences were resuspended to a concentration of 100 uM with ddH_2_O (Millipore, cat#1694). Backbones were synthesized through PCR, using a 5’ phosphorylated forward primer containing a variable input domain, and a 5’ biotinylated reverse primer containing a universal primer site (Table S1). Each 50 uL PCR reaction contained 500 nM of both forward and reverse primers, 200 uM of deoxynucleotide (dNTP) solution mix (New England Biolabs (NEB), cat# N0447L), 12 pM template, 1x ThermoPol Reaction Buffer, and 20 units/mL of Vent® (exo-) DNA Polymerase (NEB, cat# M0257L) or alternatively with Taq (NEB, cat# M0273S) and respective buffer. Reactions were placed into PCR strips at 50 uL per tube and placed into the ProFlex™ 3 x 32-well PCR System (Applied Biosystems, cat# 4484073). The PCR protocol was as follows: initial denaturation at 80℃ for 2 minutes, followed by 30 cycles of 15 seconds each of denaturing at 95℃, annealing at 56℃, and extension at 72℃. Once the protocol has finished, the reaction was allowed to extend for an additional 5 minutes before holding at 4℃ until used. All reactions were purified using a MinElute PCR Purification Kit (Qiagen, cat# 28004), using 1 column per 100 uL of PCR reaction, and eluting with 10 uL. 2 uL of each PCR reaction was used to measure its absorbance on a UV-Vis spectrophotometer: NanoDrop™ (ThermoFisher, cat# ND-ONE-W). In a final optional step, we amplified the template again using shorter (20 nt) versions of the 5’ modified primers to reduce the amount of 5’ truncated products, and therefore maximize labelling on the amplicon for better digestion efficiency at the next step.

Following amplification, dsDNA amplicons were digested with lambda exonuclease (NEB, cat# M0262S) for 10 minutes at 37℃ according to standard protocol (5 ug dsDNA / 50 uL reaction). All reactions were again purified using a MinElute PCR Purification Kit and their concentrations were adjusted to 500 nM.

### Output hybridization onto neuron backbones

ssDNA backbones (60 nM final) were mixed with ssDNA output oligonucleotides of interest at 2x excess (Table S1) in 1x annealing buffer (5 mM Tris, 1 mM EDTA, 5 mM MgCl, Millipore, cat# 648311-1KG, Bio Basic, cat# SD8135, Bio Basic, cat# MRB0328) and placed into the thermocycler. The annealing protocol was as follows: heating to 75℃ and lowering to 4℃, over the course of 2 minutes, holding at 4℃ until used.

### Solid-phase immobilization of biomolecular neuron backbone

Dynabeads™ MyOne™ Streptavidin C1 (Invitrogen™, cat# 65001) were briefly vortexed to ensure homogeneity, and suspended in 2X Binding & Wash Buffer (BWB) (10 mM Tris-HCl (pH 7.5), 1 mM EDTA, 2 M NaCl, Bio Basic, cat# DB0483). Then the solution is washed twice with 2x BWB, allowing 30 seconds for beads to magnetize between washes in the DynaMag™-96 Side Magnet (Invitrogen, cat# 12331D), and then left pelleted. A 100 uL reaction was created in the same tube with the beads, containing 30 nM annealed neuron and 1x BWB. The reaction was flicked to mix and allowed to incubate at 25℃ for 15 minutes. After this, the beads were washed with in 1x BWB heated to 40℃ four times, and left in 1X Isothermal Amplification Buffer II (NEB, cat# B0374S) until used.

In an alternate procedure, annealed neurons were immobilized onto a streptavidin coated plate (Thermo Scientific™ Pierce™, cat# 15506). Each well contained 25 uL of 30 nM annealed neuron and 1x BWB, and was allowed to incubate at 25℃ for 30 minutes. After this, wells were washed with in 1x BWB heated to 40℃, four separate times, and left in 1X Isothermal Amplification Buffer II until used.

### Polyacrylamide Gel Electrophoresis

14% denaturing polyacrylamide gels were casted in 1 mm cassettes (Invitrogen, cat# NC2010) with 10 mL solution containing 0.5X TBE Buffer (Bioshop, cat# TBE444.4), 14% Acrylamide/Bis Solution 19:1 (Bio-Rad, cat# 1610144), 2.8 g Urea (BioShop, cat# URE002.1), 0.05% TEMED (Bioshop, cat# TEM001.25), 0.1% Ammonium persulphate (Sigma, cat# A3678-100G), and allowed to cure for 2+ hours. Leftmost lane was loaded with Ultra Low Range DNA Ladder (Invitrogen, cat# 10597012), and samples were mixed 1:1 with a denaturing dye containing 95% formamide (Sigma, cat# F9037-100ML), 0.02% sodium dodecyl sulfate (SDS) (Bioshop, cat# SDS001.1), 1 mM EDTA (Sigma, cat# 324504-500ML) and bromophenol blue (Sigma, B0126-25G). Gels were developed for 20 minutes at 270V, and subsequently stained with Diamond™ Nucleic Acid Dye (Promega, H1181) for 10 minutes. Images were captured using the iBright FL1500 Imaging System (Invitrogen, cat# A44241) using auto exposure and analyzed using Fiji^45^.

### Polymerase-mediated strand displacement synthesis reactions

Twenty-four uL of output-hybridized biomolecular neurons were placed into a PCR strip, and the supernatant was removed. For each reaction, a 12.5 uL of 2x master mix was created containing 2X Isothermal Amplification Buffer II, 600 uM dNTP mix, 250 U/mL Bst 3.0 DNA Polymerase (NEB, cat# M0374S), 2.25 ng/uL Single-Stranded DNA Binding Protein (Sigma, cat# S3917 OR Promega, cat# M301A). 12.5 uL of master mix was mixed with 12.5 uL of 2x input oligonucleotide, and 24 uL of this mixture was placed into supernatant removed beads. The mixture was mixed well to ensure homogeneity and 22 uL was pipetted into a black, flat-bottom 384-well plate (Greiner Bio-One, cat# 784900) and placed into a monochromator-based plate reader heated to 40℃: Synergy H1 Microplate Reader (BioTek Instruments, cat# 8041000), reading every 20 seconds at excitation/emission: 488/517 nm. Clear plate-sealing films were found to introduce high background while reducing signal, and thus were not used for short read times (∼15 - 30 minutes). A similar protocol was used when immobilized on streptavidin-coated plates instead.

## ACKNOWLEDGEMENTS

R.C.L acknowledges individuals and organizations who contributed vector graphics to bioicons.com under CC0 or CC-BY 3.0, and the GeoDataViz Toolkit under OGL-UK-3.0, as well as the Mitacs Globalink Research Award Abroad. A.C acknowledges the NSERC Undergraduate Student Research Award. C.Y.T. acknowledges SCACE Graduate Fellowship, Kristi Piia Callum Memorial Fellowship, and Paul Starita Graduate Student Fellowships for their support. L.Y.T.C acknowledges the NSERC Discovery Grant [RGPIN-2020-05966], New Frontiers in Research Fund [NFRFE-2018-01444], CFREF Medicine by Design Initiative for funding support. We thank the members of the Chou Lab for their support and comments on this manuscript.

## AUTHOR CONTRIBUTIONS

R.C.L and L.Y.T.C conceptualized the project and designed the experiments. R.C.L performed the experiments with assistance from A.C and C.Y.T. R.C.L performed analyses on the data. C.Y.T and L.Y.T.C participated in data interpretation and provided suggestions. L.Y.T.C supervised the project. R.C.L and L.Y.T.C wrote the manuscript with input from all authors.

## SUPPLEMENTARY INFORMATION

**Supplementary Figure 1.**
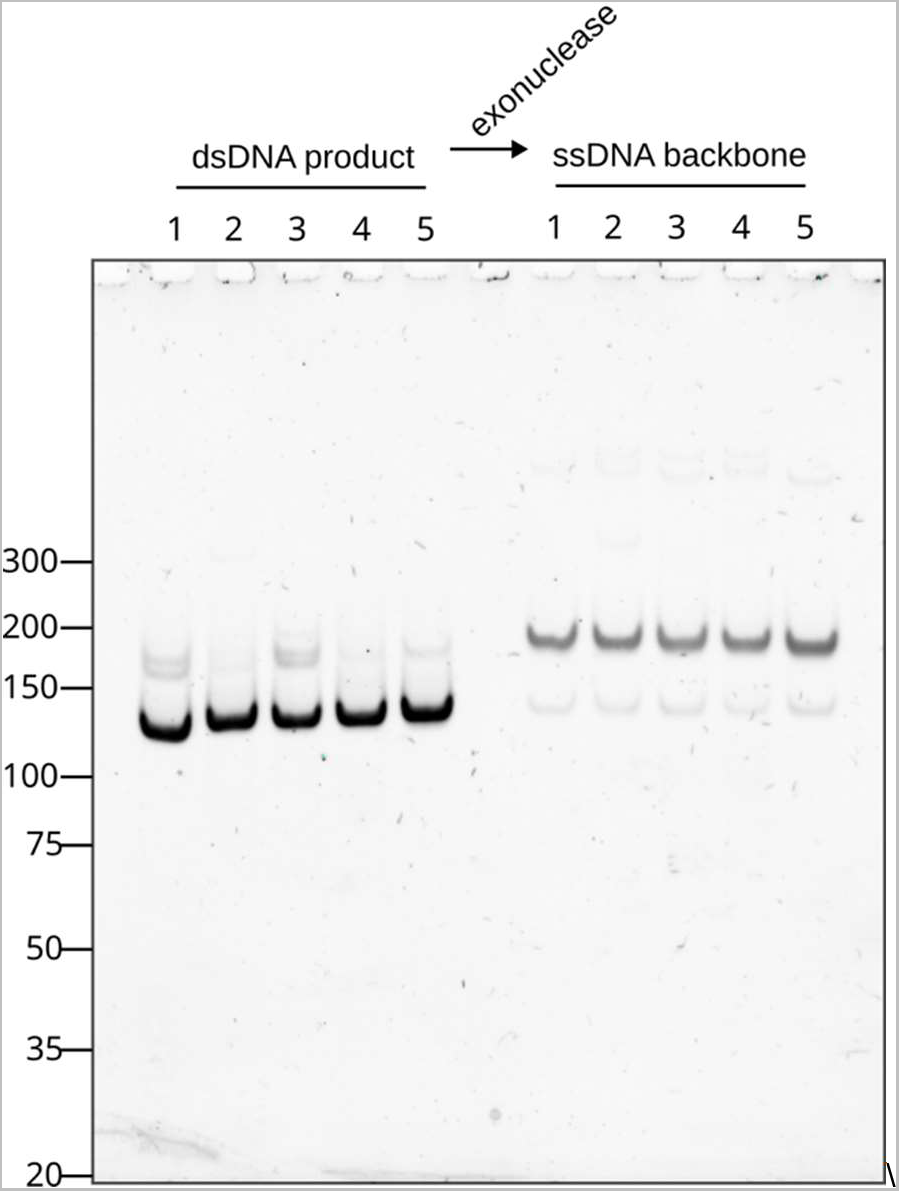
Enzymatic synthesis enables appending of a variable xn domain to universally addressed backbone. Five 5’ phosphate modified forward primers (43 nt), one 5’ biotin modified primer (20 nt), and one template with 5 universal addresses (112 nt) (Supplementary Table 1) were used in PCR to create five backbones. **(Left)** 8% Native PAGE shows the five distinct 135 nt dsDNA products. **(Right)** Lambda exonuclease degrades phos-modified DNA in dsDNA complex to yield ssDNA, with 4.8 ± 0.6% of dsDNA remaining (mean ± SEM, n = 5). **Note:** While the synthesis of 5 templates were shown in the data, only 3 were used in this work (1, 2, 5).

**Supplementary Figure 2.**
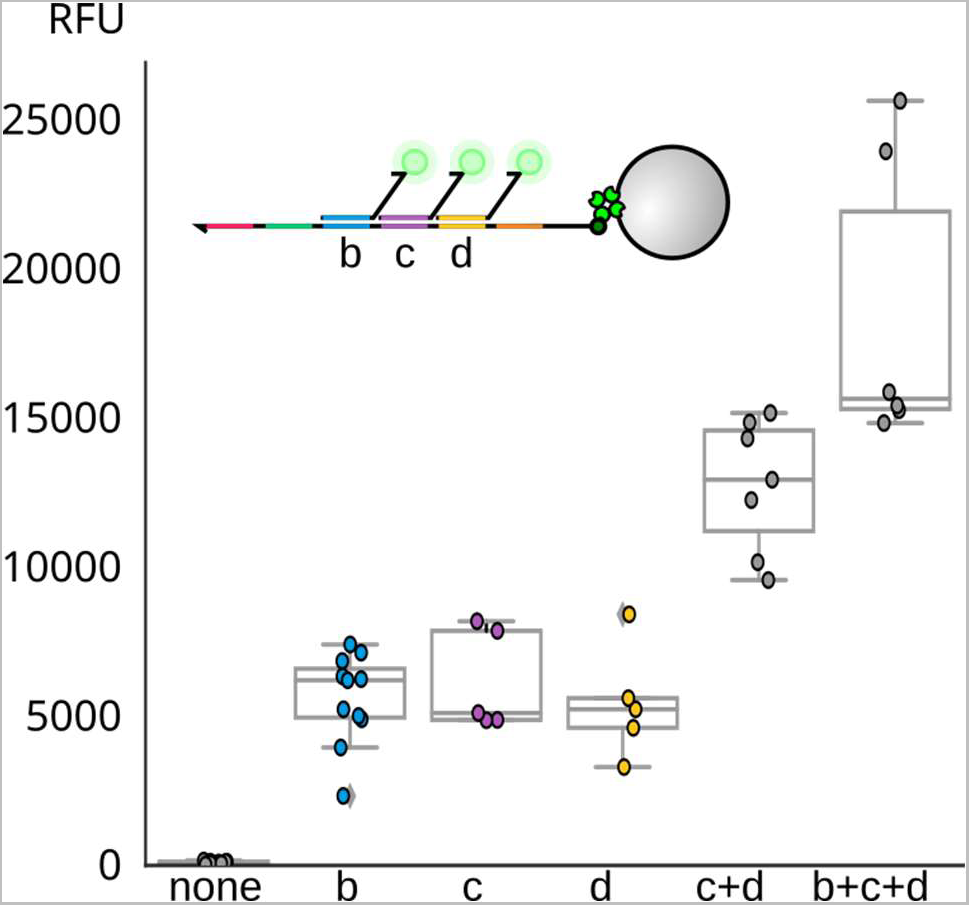
Large batch-to-batch variability of on-surface ssDNA with NaOH denaturation. Neurons synthesized through a NaOH denaturation method were labelled with FAM-fluorescent outputs in addresses b,c, and d. After washes, fluorescence of each sample was measured using a spectrophotometer (n = 6). Data suggests that neurons synthesized this way exhibit large variation in the amount of ssDNA retained on the bead surface.

**Supplementary Figure 3.**
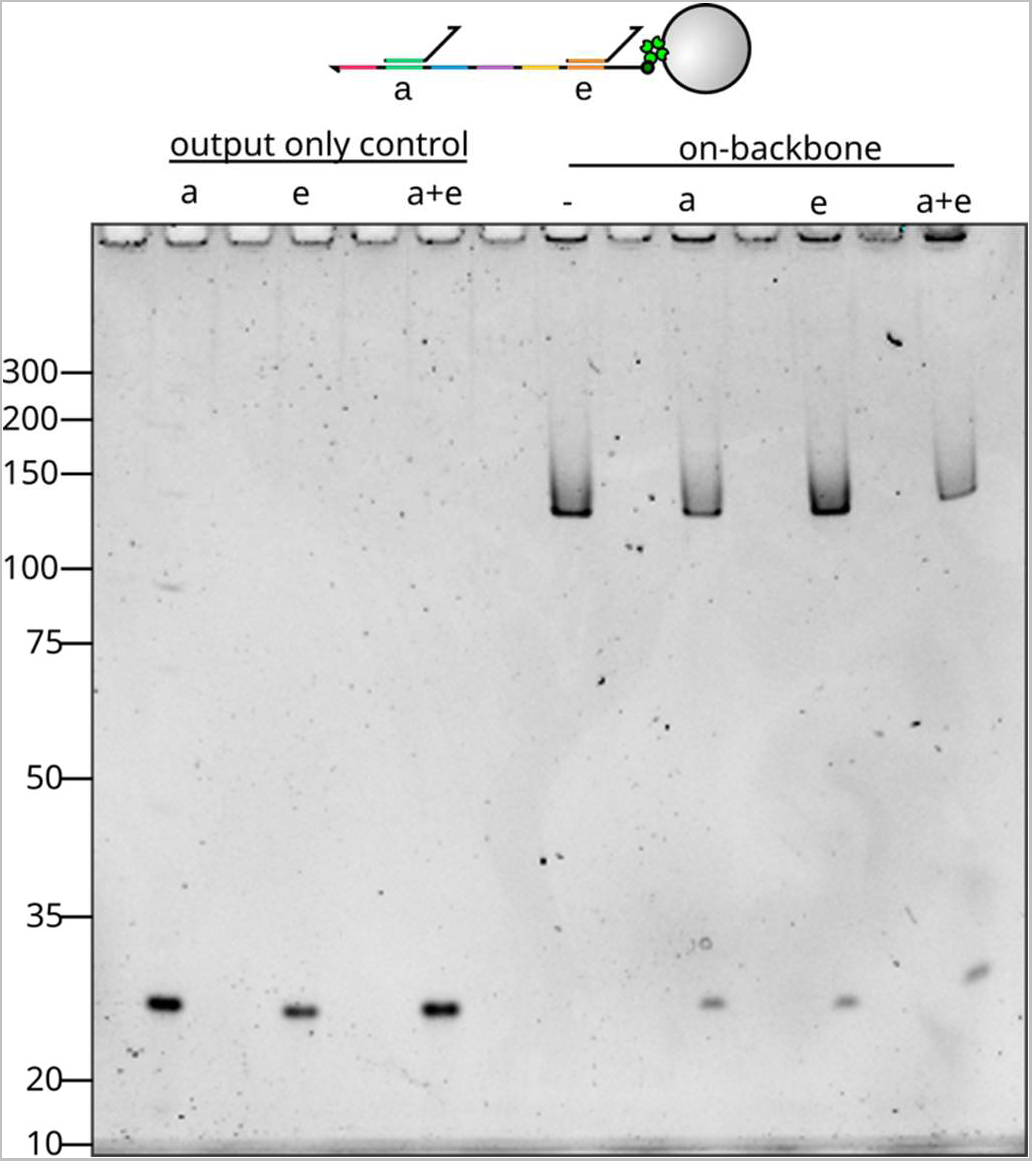
Addressability of neuron backbone allows for decoration using ssDNA outputs. Hybridization of output strands to neuron backbone on the two addresses most proximal (a) and distal (e) to the input binding region. PAGE (14%, denaturing) results show that output strands can be successfully placed addresses a and e of ssDNA backbone. Together with Figure 2b, results show complete addressability of neuron backbone without positional dependencies.

**Supplementary Figure 4.**
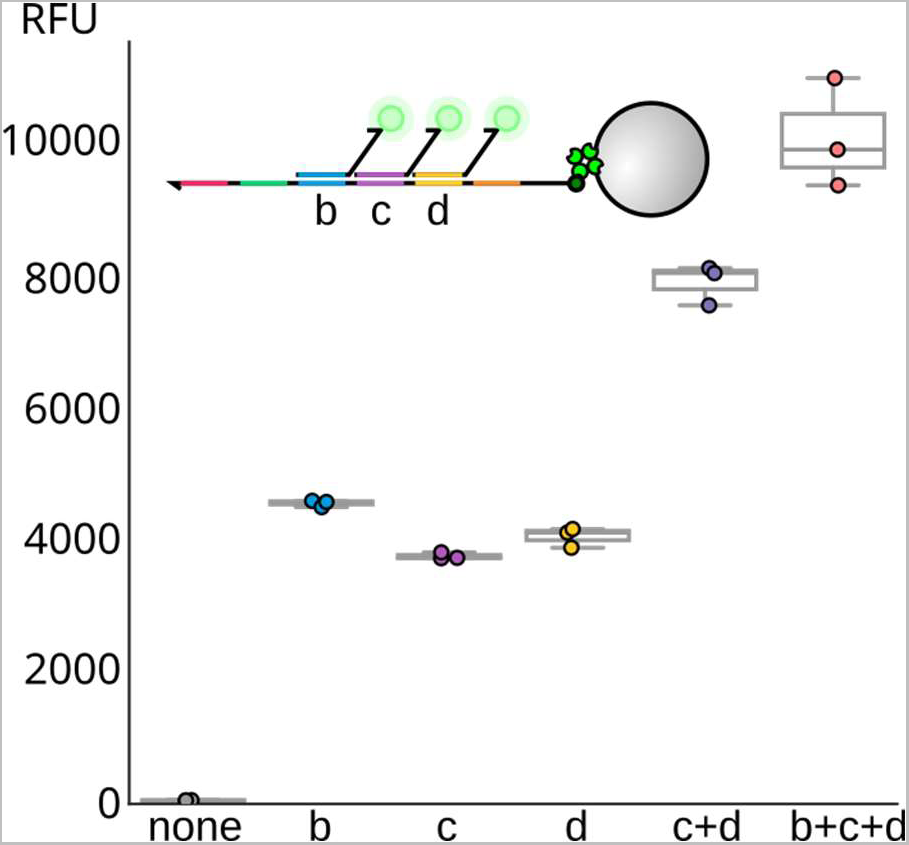
Fluorescent labelling of ssDNA neuron backbone. ssDNA outputs with 3’ FAM modifications were annealed onto ssDNA neuron backbones in different combinations, and fluorescence was observed using a spectrophotometer. We observed no significant differences between single-output backbones, while two- and three-labelled outputs showed 2x and 3x quantized increases in fluorescence. These results suggest complete labelling of address without competitive inhibition.

**Supplementary Figure 5.**
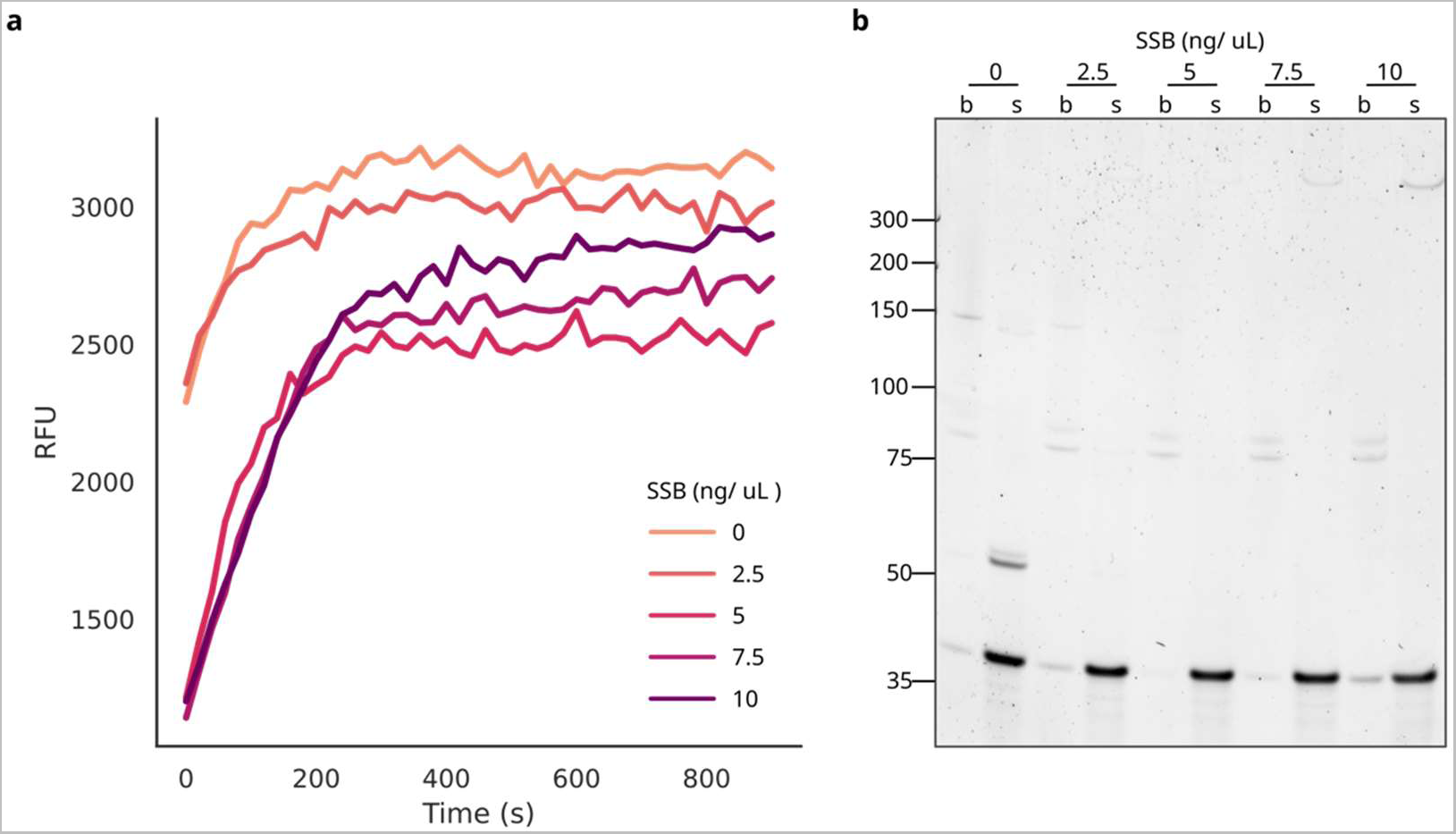
Single-Stranded Binding Protein (SSB) reduces formation of non-specific products, but slows reaction kinetics at high concentrations. **a**, Kinetic data of primer extension assay with increasing amounts of single-stranded binding protein (SSB) in reaction. Data shows that SSB slows reaction and reduces maximum fluorescent signal at concentrations greater than 2.5 ng/uL. **b,** Primer extension assay with single-stranded binding protein (SSB). Lanes represent bead solutions (b) and reaction supernatants (s) at increasing concentrations of SSB. Bead solutions contain extended dsDNA products, while supernatants contain displaced output strands. Denaturing PAGE results show that the SSB >= 2.5 ng/uL reaction prevents synthesis of non-specific DNA products indicated by the ∼55 nt band in the 0 ng/uL s condition. Together, kinetic and gel data suggest that SSB concentration at 2.5 ng/uL will prevent the formation of non-specific products without slowing reaction kinetics compared to a no SSB control. No fluorescence normalization was performed on this data.

**Supplementary Figure 6.**
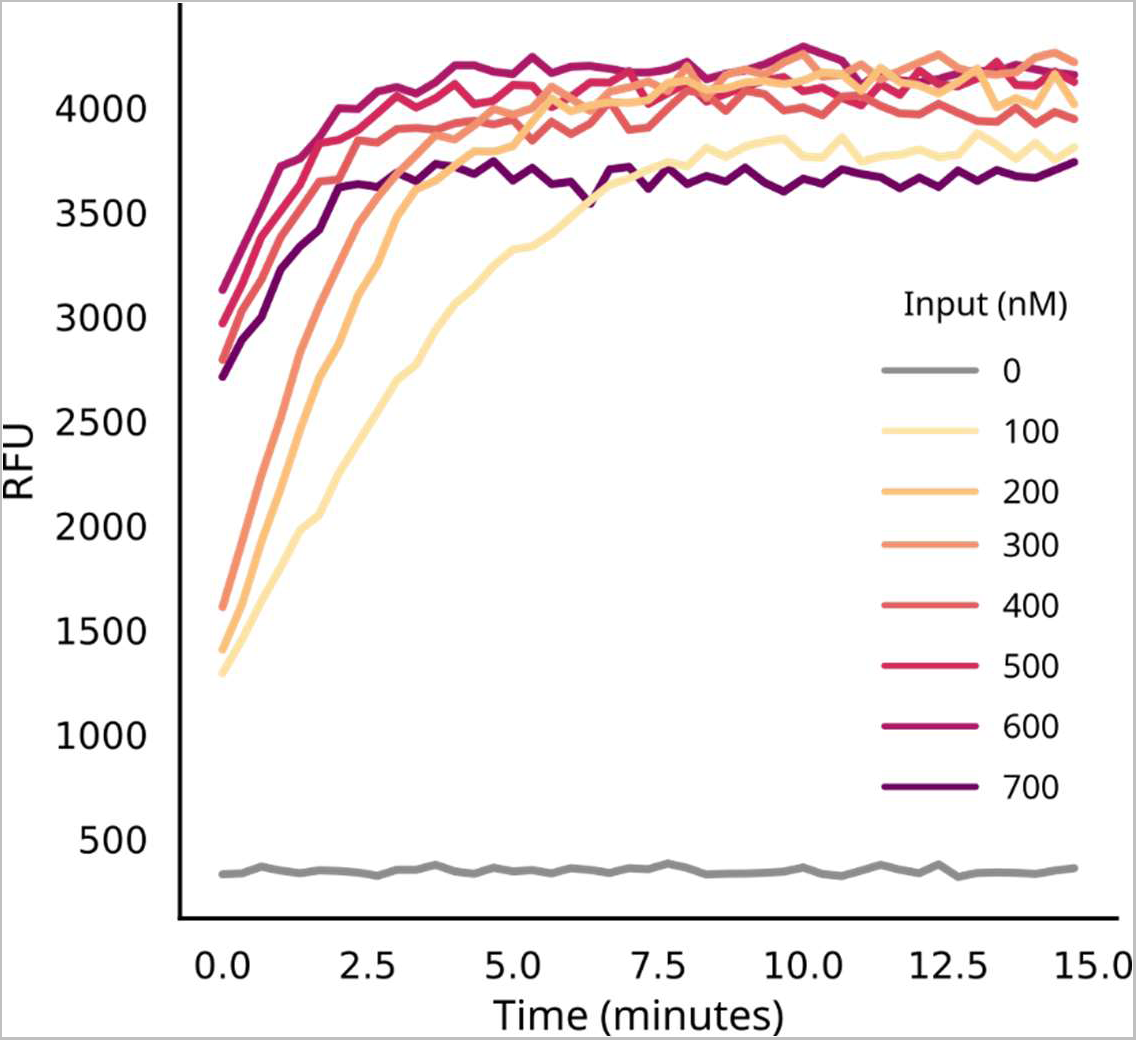
Kinetic response of biomolecular neurons to large concentrations of input signal (>100 nM). 1-input 2-address backbones were functionalized with fluorophore quencher (BHQ2/FAM) pairs and reacted with input concentrations ranging from 100 nM to 700 nM. Data suggests that all concentrations above 100 nM achieve the same maximum signal, with larger concentrations reacting more quickly. No fluorescence normalization was performed on the data.

**Supplementary Figure 7.**
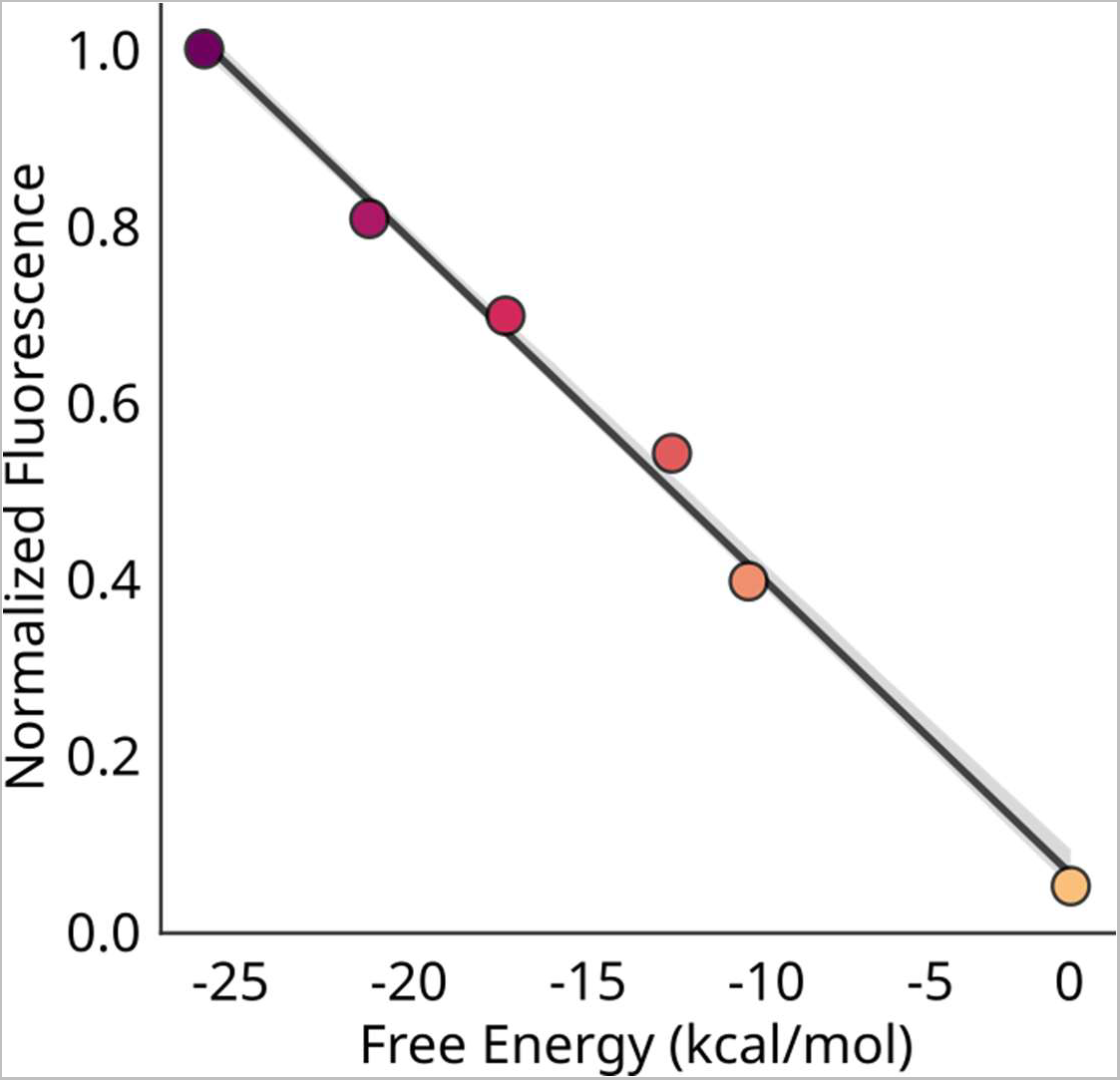
Kinetic response of biomolecular neurons to changes in input hybridization energy. Data from Figure 3d plotted against hybridization energy instead of nucleotide length. Results show a linear relationship between hybridization energy and neuron activation.

**Supplementary Table 1.**
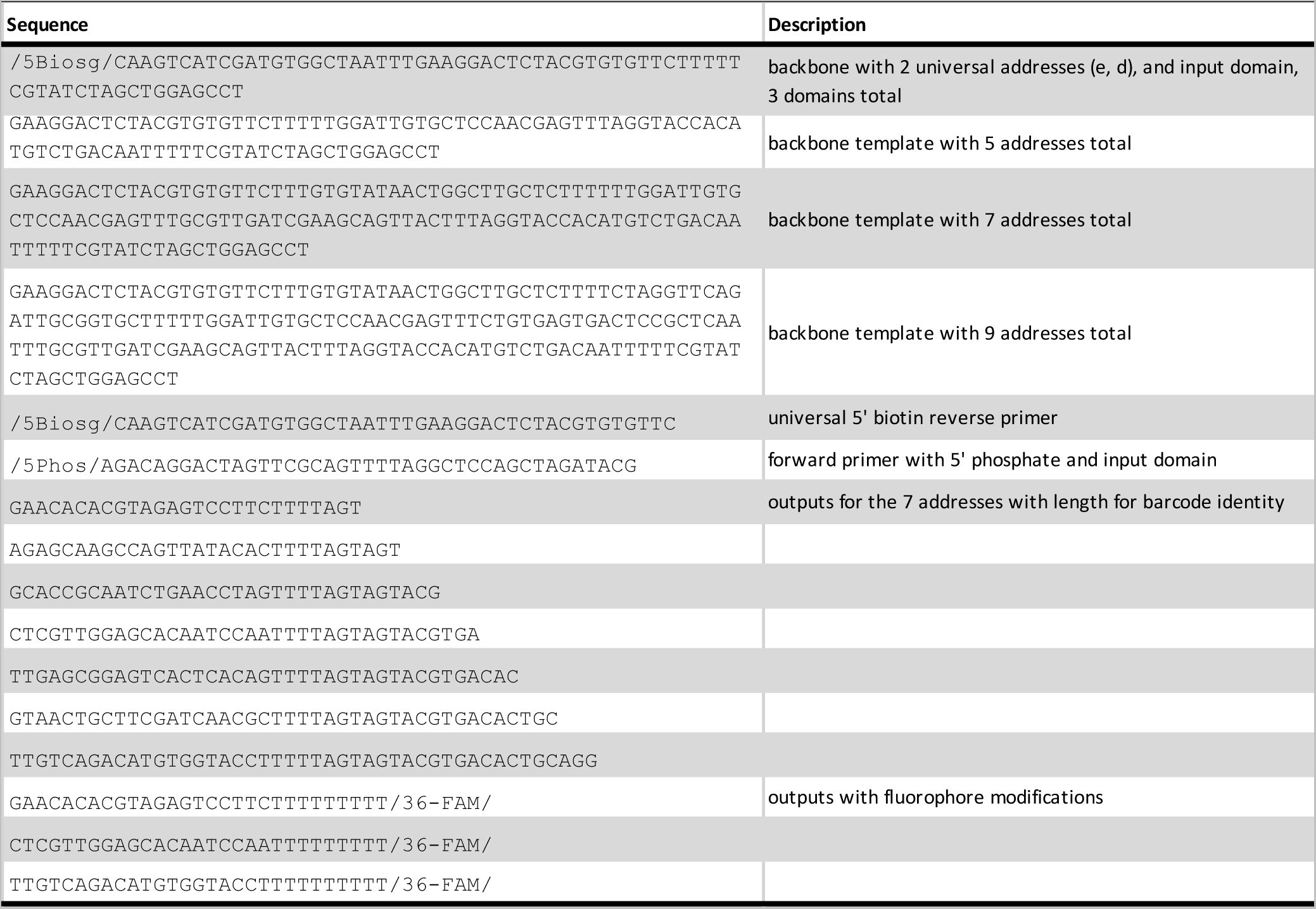
List of DNA sequences used in for synthesis verification in Figure 2. Table contains a list of sequences used to create the neurons for dPAGE visualization, as well as descriptions of the purpose of each sequence. Sequences were designed using Python and NuPACK 4.0.

**Supplementary Table 2.**
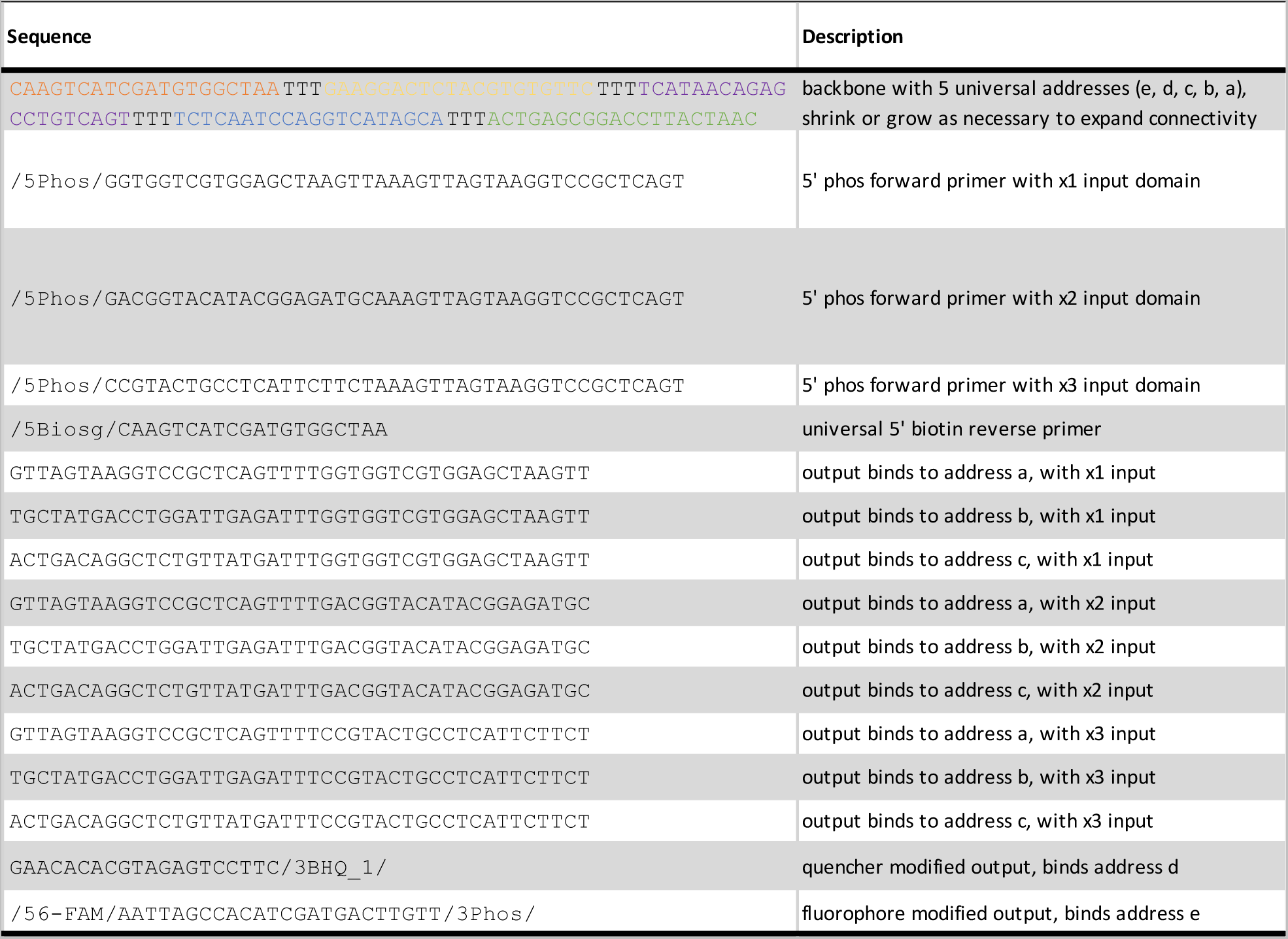
List of DNA sequences used in for synthesis verification in Figure 3-4. Table contains a list of sequences used to create the neurons for kinetic experiments, as well as descriptions of the purpose of each sequence. Sequences were designed using Python and NuPACK 4.0.

**Supplementary Table 3.**
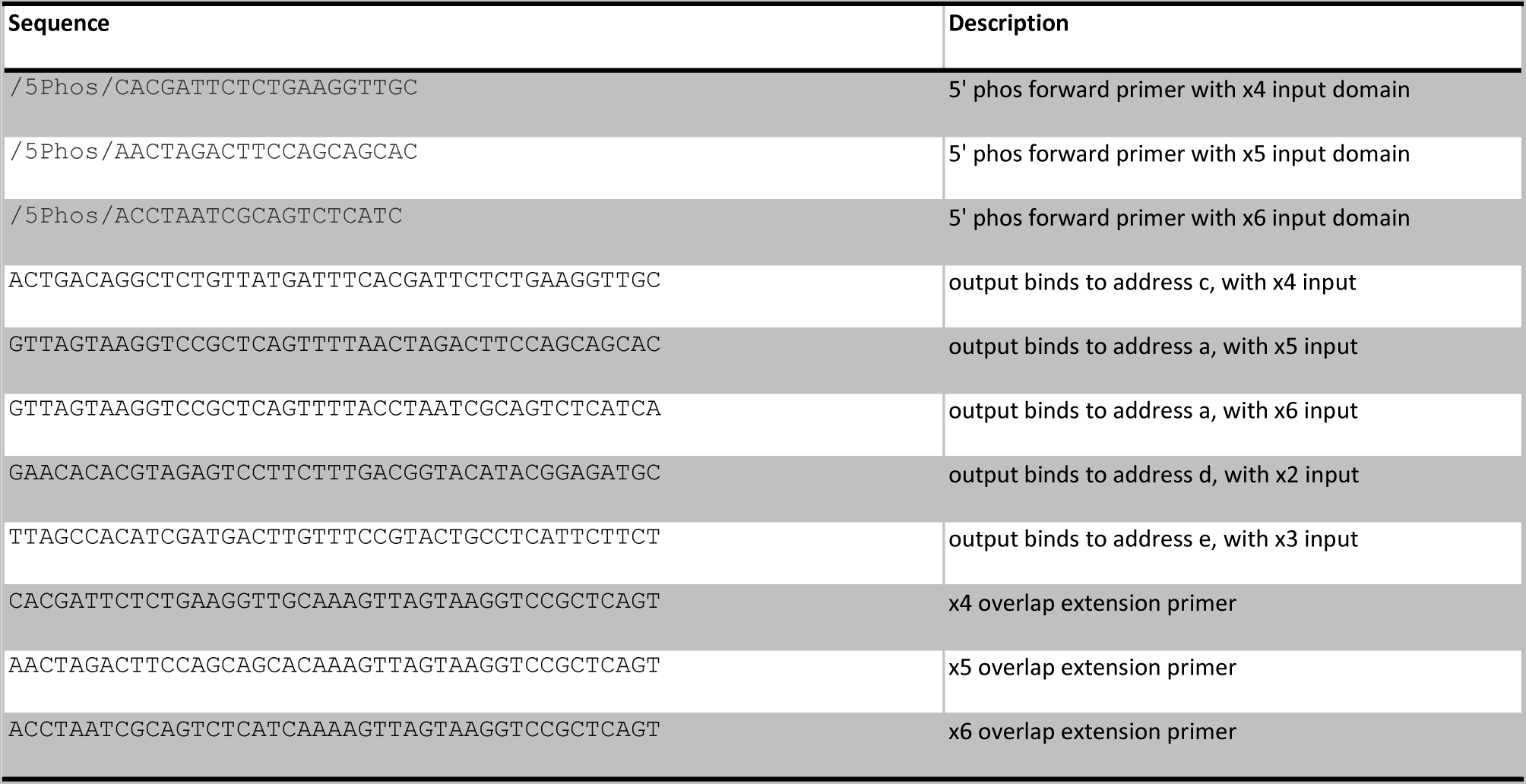
List of DNA sequences used in Figure 5. Table contains a list of sequences used to create three additional neurons, n4, n5 and n6, as well as descriptions of the purpose of each sequence. Shorter 5’ phos primers were used to ensure better modification efficiency, and longer unlabelled overlap extension primers Sequences were designed using Python and NuPACK 4.0.

